# Five new *Caenorhabditis* species from Indonesia provide exceptions to Haldane’s rule and partial fertility of interspecific hybrids

**DOI:** 10.1101/2025.05.14.653126

**Authors:** Mia Prastika Devi, Elkana Haryoso, Emha Ilhami Rais, Anggik Karuniawan, Minhajul Qowim Yahya, Aurélien Richaud, John Wang, Matthew V. Rockman, Hagus Tarno, Marie-Anne Félix

## Abstract

Given the interest in the biogeography and diversity of the *Caenorhabditis* genus, we established a collection of these nematodes from field surveys on four Indonesian islands. We isolated over 60 *Caenorhabditis* strains belonging to ten species. Five species were previously known from other locations: *C. briggsae*, which was predominant, *C. tropicalis, C. nigoni, C. brenneri* and *C. elegans*. The five other species are new, and we describe them here as *Caenorhabditis indonesiana, Caenorhabditis malino, Caenorhabditis ceno, Caenorhabditis brawijaya* and *Caenorhabditis ubi*. RNA sequence analysis of 1,861 orthologous genes placed all species from Indonesia in the *Elegans* group of *Caenorhabditis* species. Four of the new species belong to a *Sinica* subclade of species so far only found in an East Asia-Indo-Pacific world region. The fifth new species, *C. indonesiana*, appears as the sister of the *C. tropicalis*-*C. wallacei* pair, both also found in Indonesia. The present findings are thus consistent with diversification in the *Elegans* group having occurred in this world region. Crosses between closely related species showed counterexamples to Haldane’s “rule”: for two pairs of species, in one cross direction we only found hybrid males. In addition, we found a pair of species that could partially interbreed: *Caenorhabditis ubi* (East Java) with *C*. sp. 41 (Solomon islands), with the hybrid males in one cross direction being fertile. Such closesly related species pairs are good models for genetic studies of incompatibilities arising during speciation.

**Summary:** This work addresses their biodiversity, phylogenetic relationships and genetic incompatibilities of *Caenorhabditis* nematodes, which are laboratory model organisms. Through field studies, the authors isolated 60 *Caenorhabditis* strains in Indonesia, representing ten species, including five new. From RNA sequencing and phylogenetic reconstruction, all ten species belong to the *Elegans* group of *Caenorhabditis*. In crosses between closely related species, the hybrid progeny can be all females, abiding by Haldane’s rule, but in other cases all males. In one species pair, partially fertile hybrids are produced in one cross direction. These closely related species are good models for studying genetic incompatibilities.

## Introduction

*Caenorhabditis elegans* is a free-living nematode species that is widely used as a biological model organism. Building a solid evolutionary framework of genetic and phenotypic variation within and around this model species is thus important for providing a context for its biology and as resource for evolutionary biology and ecology. The last 20 years have seen a blossom of discovery of new *Caenorhabditis* species (Kiontke et al. 2011; Félix et al. 2014; Huang et al. 2014; Ferrari et al. 2017; Slos et al. 2017; Kanzaki et al. 2018; Crombie et al. 2019; Stevens et al. 2019; Sloat et al. 2022). Many species including *C. elegans* can be found in rotting vegetal material rich in bacteria, such as decomposing fruits, flowers and stems (Kiontke et al. 2011; Schulenburg and Félix 2017). These species, especially their larval diapause stage called the dauer larva, can be carried between food patches by larger invertebrates such as mollusks and arthropods (Félix and Braendle 2010).

The present phylogeny and biogeography of the *Caenorhabditis* genus indicate that the *Elegans* group of species may have radiated in the Asia-Pacific region (Frézal and Félix 2015; Stevens et al. 2019). The most divergent populations of *C. elegans* were found in the Hawaiian islands and the rim of the Pacific ocean (Crombie et al. 2019; Lee et al. 2021; Crombie et al. 2023). The present sister species of *C. elegans*, named *C. inopinata*, was found in South Japan on tropical figs and their associated wasps (Kanzaki et al. 2018; Woodruff and Phillips 2018). In Indonesia, another fig-associated species provisionally called *Caenorhabditis* sp. 35 was found in West Sumatra (Jauharlina et al. 2022), and an *Elegans* group species, *C. wallacei*, was described from Bali (Kiontke et al. 2011; Félix et al. 2014). The sampling of *Caenorhabditis* strains from Indonesia was thus likely to yield new species, as well as insights about their habitat and diversity.

Most *Caenorhabditis* species reproduce through males and females. Only three species, *C. elegans, C. briggsae* and *C. tropicalis*, independently evolved a mode of reproduction via selfing hermaphrodites and facultative males (Kiontke et al. 2011). Finding a closer sister species to *C. elegans* would be valuable. In addition, pairs of species that are partially cross-fertile, such as *C. briggsae*-*C. nigoni* (Woodruff et al. 2010; Bi et al. 2015; Bundus et al. 2015) or *C. latens*-*C. remanei* (Dey et al. 2012), provide good cases for studies of genetic incompatibilities arising during speciation (Cutter 2017).

Given the interest in the biogeography of the *Caenorhabditis* genus, the quest for closely related species to study speciation mechanisms and the paucity of previous sampling, we aimed to collect and culture *Caenorhabditis* from Indonesia. Diverse elevations and landscape types (e.g. forests, agricultural land) and various sample types (rotting vegetal and fungal matter, such as flowers, fruits, stems, leaves, wood, fungi) were collected. We found eleven different species and could maintain ten in culture, five of which are new species. All are in the *Elegans* group of *Caenorhabditis* species. Interestingly, crosses between closely related species showed only yielded F1 hybrid males, providing counterexamples to Haldane’s rule. In addition, we found a pair of species (*C. ubi* n. sp. and *C*. sp. 41 from the Solomon islands) that could partially interbreed.

## Materials and Methods

### Sampling and isolation

Using a field survey method, locations were selected based on suitability with the habitat of nematodes of the genus *Caenorhabditis*. Sampling was conducted across several regions in Indonesia, at various points on Java, Sulawesi, Lombok and Bali islands. Each observation location was marked using a Global Positioning System (GPS) to record the altitude and coordinates of the observation points. On Java Island, samples were collected from Coban Talun in Wonorejo Hamlet, Tulungrejo Village, Bumiaji District, Batu City; University of Brawijaya Forest in Tawang Agro Village, Karangploso Sub-district, Malang Regency; and Mount Bromo in Ngadas Village, Sukapura Sub-district, Probolinggo Regency. On Sulawesi Island, sampling was carried out in two villages in Gowa Regency, Central Sulawesi Province, Malino and Lanna Villages. From Lombok Island, West Nusa Tenggara Province, samples were collected from two villages: Lingsar Village in West Lombok Regency and Setiling Village in Central Lombok Regency. From Bali Island, samples were collected in Sayan, Ubud, Megwi, Marga, Ababi and Besakih. The extraction and identification processes were conducted at the Laboratory of Plant Pathology I and Laboratory of Plant Pest II, Department of Plant Pests and Diseases, Faculty of Agriculture, Universitas Brawijaya, Malang, East Java, Indonesia, and at the Institut de Biologie de l’Ecole Normale Supérieure (IBENS), Paris, France.

For nematode extraction and culture in Malang, we used Petri dishes containing Nematode Growth Medium (NGM) with *Escherichia coli* OP50 placed at the centre. The preparation began by making the NGM. A total of 14.5 grams of NGM was added to a 1-liter Schott bottle, followed by 500 mL of distilled water. The Schott bottle was then placed in a clear polypropylene (PP) plastic bag and autoclaved at 121°C and 20 psi pressure for an hour. After autoclaving, the bottle was allowed to cool for 10 minutes before the medium was aseptically poured into glass or plastic Petri dishes of 90 or 55 mm diameter for extraction and culture, respectively. Once the medium solidified, 100 μL of *E. coli* OP50 culture was added to the center of each dish using a micropipette. The pouring of NGM and addition of *E. coli* OP50 culture were conducted in a Laminar Air Flow Cabinet (LAFC) to prevent contamination. The *E. coli* on the NGM was incubated for 24 hours at room temperature.

Table S1 provides the time of field collection and of plating the samples in the laboratory. The plating of samples was performed in a time interval varying from one day to two weeks for samples analyzed in Paris. This study employed the Agar Culture Plate extraction method (Barrière and Félix 2014). Ca. 10 g samples of substrate were collected, with the plant samples being cut into smaller pieces using scissors. These samples were then evenly spread in a circular pattern along the edge of the NGM plate. The plates were left at room temperature to allow nematodes to migrate from the substrate toward the *E. coli* OP50 on the NGM. Each plate was monitored several times (from a few hours to a few days) and nematodes that migrated to *E. coli* were transferred aseptically using a nematode platinum wire pick to a fresh *E. coli* OP50-seeded NGM plate. The nematodes initially transferred to to establish an isofemale line were single hermaphrodites, single mated females or a pair of nematodes (male and female), ensuring that the population contained a single species.

### Culture and freezing

Isofemale strains were maintained and frozen using standard methods. The nematodes cultured on NGM plates were transferred to fresh plates either using the pick or the “Chunking” method (Stiernagle 2006). This latter method involved cutting a section of agar from the old NGM plate using a sterilized syringe or spatula and transferring the agar piece to a new NGM plate.

Nematodes were frozen in Paris with standard *C. elegans* protocols (Stiernagle 2006). All strains thrive at 25°C, except *C. elegans* HPT48, and *C. brawijaya* n. sp. HPT49 and HPT50, which grow better at 20°C.

### Mode of reproduction, crosses and assignment of strains to biological species

The mode of reproduction of each strain was assessed by isolation of L4 larvae with a female body. In these circumstances, selfing hermaphrodites produce progeny, and females fail to produce progeny.

Crosses between male-female strains were established by placing together five L4 females of one strain and five L4 to adult males of another strain. The plate was checked for the presence of cross-progeny (F1) and later cross-fertility of the F1 animals over several days. In some cases, backcrosses of the F1 animals to the parental strains were performed.

Species with selfing XX hermaphrodites produce rare X0 males that can be crossed like *C. elegans*. For crosses with these species, selfing hermaphrodites were crossed to males, also using five animals of each sex. A cross was considered positive when the progeny included a high proportion of males (>20%).

All strains and their geographic position can be found and visualized on a world map on the following website: https://justbio.com/tools/worldwideworms/search.php

### ITS2 sequencing

Sequencing of the ribosomal DNA internal transcribed spacer ITS2 was performed as in (Kiontke et al. 2011), using primers 5.8S-1 and KK28S-22 for amplification and KK28S-22 for Sanger sequencing. For the ITS2 sequences that were highly polymorphic (HPT35 and HPT43), the fragments were cloned into the pGEM-T easy vector (Promega) and sequenced with a T7 primer. The ITS2 sequences are deposited at GenBank with accession numbers PV569245 -PV569249.

### RNA sequencing

Before RNA preparation, the reference strains HPT10, HPT35, HPT5 were inbred by crossing a single L4 female and a male (brother-sister mating) for five generations. HPT43 was further inbred for 25 generations yielding inbred strain JU4643. HPT50 was used without inbreeding. RNA was prepared from a mixed-stage population using Trizol and a freeze-thaw cycle as in (Stevens et al. 2019). Sequencing libraries were prepared from polyA RNA and sequenced on an Illumina NovaSeq with 150 bp paired-end reads.

### Phylogeny

We estimated the phylogeny of the *Elegans* group from the amino-acid sequences of single-copy proteins, following the procedure in (Sloat et al. 2022). First, RNAseq read-pairs were trimmed using trimgalore 0.6.6 (https://github.com/FelixKrueger/TrimGalore) in paired mode with q=25, and transcriptomes assembled using Trinity 2.15.1 (Grabherr et al. 2011) with default settings. The longest isoform for each transcript was extracted with transdecoder 5.5.0 (https://github.com/TransDecoder/TransDecoder) and the resulting transcriptomes processed through BUSCO 5.3.0 (Seppey et al. 2019) to identify genes representing 3131 core gene models in the nematoda_odb10 database. We next collected analogous data from an additional 21 *Elegans* group species and used busco2fasta.py (https://github.com/lstevens17/busco2fasta) to collect sets of BUSCO genes present as single-copy genes in the transcriptomes at least 80% of the species. The predicted amino acid sequences of each gene were then aligned using MAFFT 7.475 (Katoh and Standley 2013) with automatic parameter selection, and the alignments trimmed to alignable positions using trimAl 1.4.1 (Capella-Gutierrez et al. 2009) with settings gt 0.8, st 0.001, resoverlap 0.75 and seqoverlap 80. We estimated the gene tree for each of the 1861 protein alignments using iqtree 1.6.12 (Nguyen et al. 2015) under the LG+I+G substitution model. We estimated the species tree from the collection of gene trees using Astral 5.7.8 (Zhang et al. 2018), and we estimated branch lengths for the species tree using iqtree -te with the concatenation of all of the protein sequence alignments (755373 amino acid positions), which we generated with catfasta2phyml (https://github.com/nylander/catfasta2phyml). Plots were generated with the ape package (Paradis and Schliep 2019). RNAseq data and assembled transcriptomes are available under NCBI BioProject PRJNA1256413.

### Morphology and Nomarski micrographs

Pictures of whole animals on the agar plates were taken using a Nikon Multizoom AZ100 equipped with a Hamamatsu Orca-flash 4.0 camera. Morphology was observed under Nomarski microsopic illumination with a 100x objective on an AxioImager 2 (Zeiss), after mounting the animals on a Noble agar pad (Shaham 2006). Pictures were taken using a Photometrics CoolSNAP ES CCD camera. Pictures showing extruded spicules were obtained after exposing the animals for 2-4 seconds in a microwave before adding the coverslip.

## Results

### Field sampling of *Caenorhabditis*

We sampled on four islands, Java, Bali, Lombok, Sulawesi (Figure 1), and collected samples at different elevations between sea level and 2400 meters. We collected samples of decomposing vegetal matter and a few invertebrates, such as mollusks, annelids and arthropods (Table S1).

**Figure 1.**
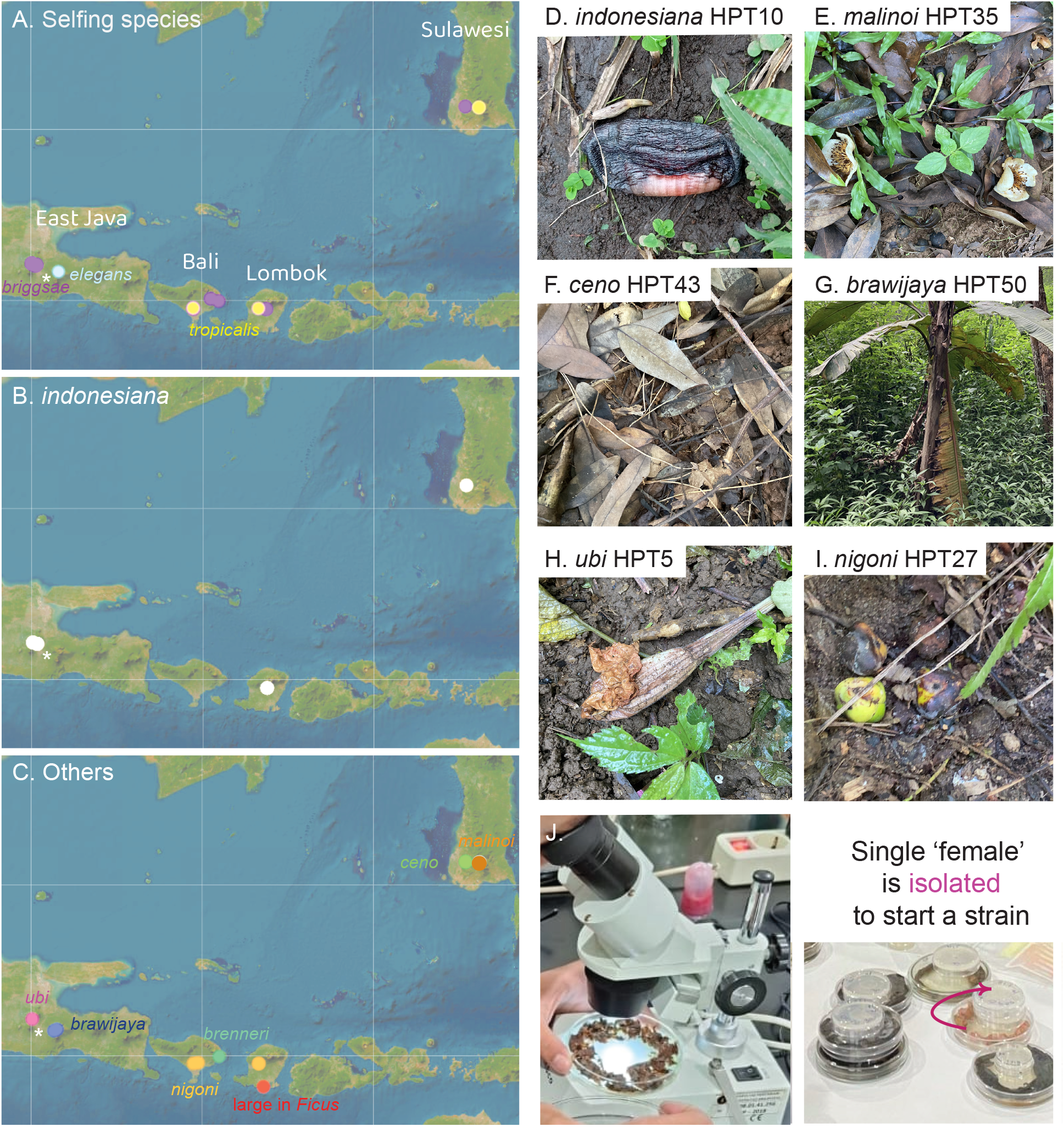
Geographic distribution of *Caenorhabditis* collected in Indonesia. (A) Geographic distribution of the three selfing *Caenorhabditis* species. *C. briggsae* is found at most sampling sites. (B) Geographic distribution of *Caenorhabditis indonesiana* (sp. 77) on several islands. (C) Geographic distribution of other species. The species are color-coded. The asterisk designates the city of Malang. Map credit: ESRI. (D-I) Pictures of representative samples. Shown are the five samples that yielded the reference strain of each new species and a rotting fruit that yielded *C. nigoni*. (D) HPT10: decomposing banana flower. (E) HPT35: *Stewartia pseudocamellia* flowers in various stages of decomposition. (F) HPT43: forest leaf litter. (G) HPT50: decomposing *Musa* pseudostem. (H) HPT5: decomposing *Brugmansia* flower. (J) Procedure for *Caenorhabditis* isolation. The samples are brought back to the laboratory and placed on agar plates (diameter 90 mm) seeded with *E. coli* bacteria. They are examined under a dissecting microscope. Single females or hermaphrodites are picked to a smaller plate (diameter 55 mm) to start an isofemale line.

Out of 204 samples collected in April-May 2024 (Table 1, Table S1), 58 were positive for *Caenorhabditis* and some samples yielded two or three *Caenorhabditis* species. From these positive samples, we established 61 isofemale strains that could be maintained in long-term culture and frozen. In addition, one sample of fresh *Ficus septica* figs yielded a large *Caenorhabditis* species that could only be maintained for few generations; these animals could be *C. inopinata* (Kanzaki et al. 2018; Woodruff and Phillips 2018) or the species isolated in (Jauharlina et al. 2022). One additional strain (HPT1) was isolated a few weeks earlier from an apple orchard. Out of the 62 strains in long-term culture, 37 reproduced by selfing of hermaphrodites and 25 through males and females.

**Table 1.** Summary of samples, strains, species, locations.

| Location | SA | PO | ST | *Cel* | *Cbr* | *Ctr* | *Cbn* | *Cni* | 77<br>*Cid* | 78<br>*Cml* | 79<br>*Cce* | 80<br>*Cbw* | 81<br>*Cub* | *Cin* or<br>sp. 35? |
| --- | --- | --- | --- | --- | --- | --- | --- | --- | --- | --- | --- | --- | --- | --- |
| Batu forest,<br>East Java | 25 | 11 | 11 | - | 8 | - | - | - | 2 | - | - | - | 1 | - |
| UB Forest,<br>East Java | 41 | 8 | 10 | - | 6 | - | - | - | 4 | - | - | - | - | - |
| Bromo area,<br>East Java | 18 | 3 | 3 | 1 | 0 | - | - | - | - | - | - | 2 | - | - |
| Lombok | 46 | 10 | 10 | - | 4 | 1 | - | 2 | 2 | - | - | - | - | 1 |
| South<br>Sulawesi | 40 | 13 | 15 | - | 4 | 2 | - | - | 1 | 5 | 3 | - | - | - |
| Bali | 34 | 13 | 13 | - | 9 | 1 | 1 | 2 | - | 0 | - | - | - | - |
| **TOTAL** | 204 | 58 | 62 | 1 | 31 | 4 | 1 | 4 | 9 | 5 | 3 | 2 | 1 | (1) |

### Selfing *Caenorhabditis* species

From the results of our crosses, all selfing strains belonged to one of the three known selfing *Caenorhabditis* species. Specifically, the majority (32/37) were *C. briggsae* and this was true on all sampled islands (Figure 1, Table 1). Four strains belonged to *C. tropicalis*, found in Lombok, South Sulawesi and Bali, but this species was absent from on the East Java sites, possibly because of their higher elevation (Table S1). Finally, a single *C. elegans* isolate was found, in a sample collected at an elevation of 2,150 meters next to the Bromo volcano in East Java, in a volcanic ash landscape with a few low bushes (Figure S1).

### Male-female *Caenorhabditis* species

We first crossed the male-female *Caenorhabditis* strains from Indonesia to each other to group them into cross-compatible biological species. This defined seven biological species (Table S2). Crosses of representative strains of each group to *C. wallacei*, which had been previously isolated from Bali, were all negative. We then sequenced a PCR product corresponding to the internal transcribed sequence 2 (ITS2) of ribosomal DNA. Based on high ITS2 sequence similarity, we set up crosses for two of them with *C. nigoni* or *C. brenneri*, respectively, and both crosses were successful (Table S2). Both *C. nigoni* and *C. brenneri* are cosmopolitan in tropical areas of the world. In the present collection, we found one *C. brenneri* in Bali and four *C. nigoni* strains, two in Lombok and two in Bali (Table 1). *C. nigoni* is partially cross-fertile with *C. briggsae* (Woodruff et al. 2010; Bi et al. 2015; Bundus et al. 2015) and we note that both could be found in the same sampling locations (Table S1).

The five other species appeared novel from the ITS2 sequence. All of them were closest to species in the *Elegans* group in the clade including *C. briggsae* and *C. brenneri* but not *C. elegans*. We sequenced RNA from one representative strain of each of these five new species and assembled transcriptomes from the reads. Using protein sequences of 1861 single-copy genes for 26 species within the *Elegans* group, we estimated the phylogenetic position of the Indonesian species (Kiontke et al. 2011; Stevens et al. 2019) (Table S3, Figure 2).

**Figure 2.**
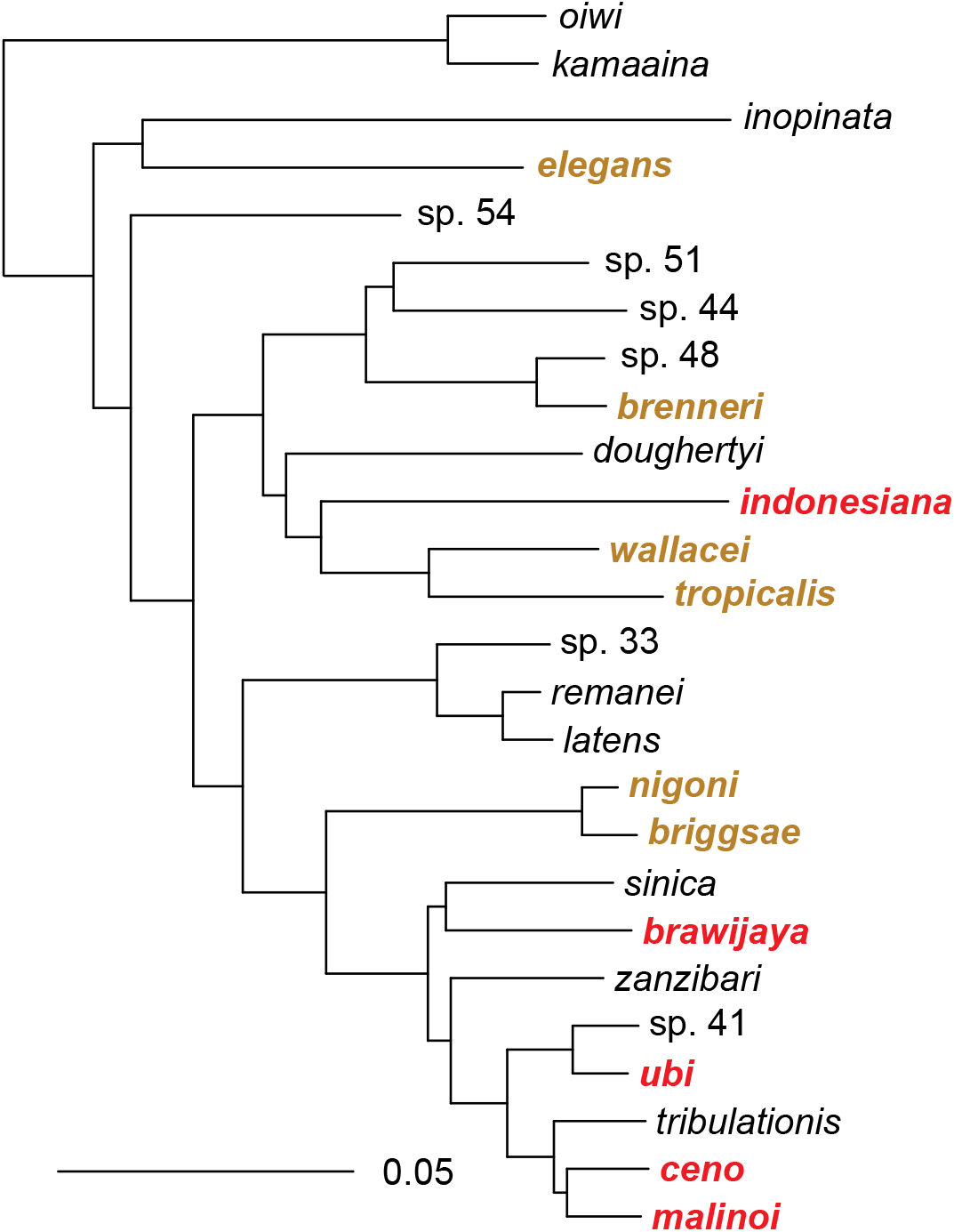
Phylogenetic relationships of the *Elegans* group of *Caenorhabditis* species. New species are in red, other species found in this study or previously in Indonesia are in brown. The topology was estimated under the multi-species coalescent from 1861 maximum-likelihood gene trees, and is shown rooted with the outgroup species *C. oiwi* and *C. kamaaina*. The scale bar represents 0.05 amino-acid replacements per site. Quadripartition support values are 1 for all branches.

One of them, represented by strain HPT10, was present on several islands, matched *C. wallacei* most closely (Table S4) but could not interbreed with it and thus defined a new species. The four others were each restricted to one island in our collection and were in a subclade including *C. sinica, C. zanzibari, C. tribulationis* and *C*. sp. 41 (represented by strain BRC20276). We crossed representative strains of each group to described species that were closest to them. Some crosses yielded larval or adult progeny but did not produce a normally fertile brood (Figure 3, Table S2). We thus raise here five new species.

**Figure 3.**
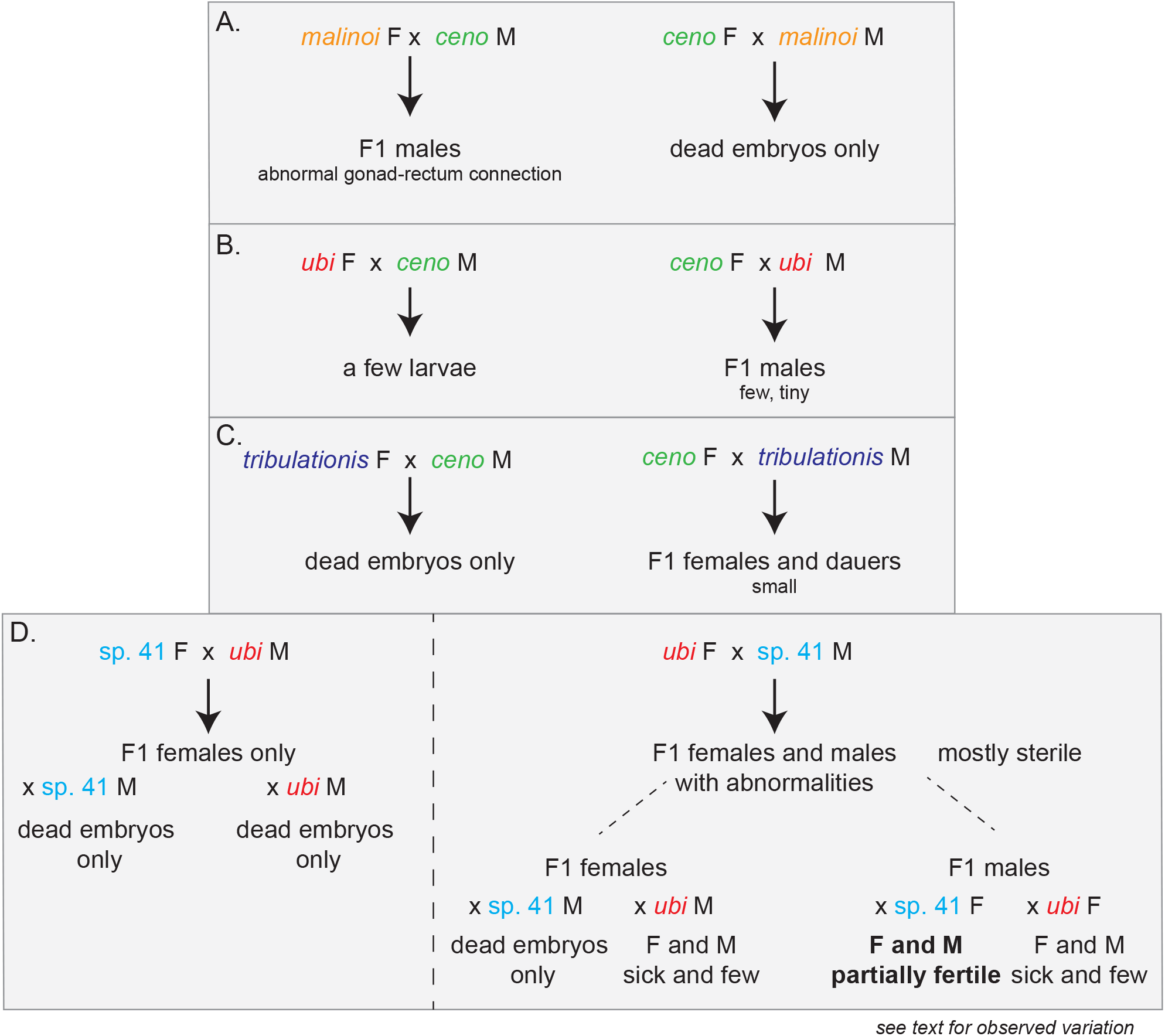
Noteworthy crosses between *Caenorhabditis* species. Panels A-C show crosses that yielded only sterile progeny, while panel D shows the crosses involving the two species that can yield some F2 (and further) progeny. Corresponding pictures are shown in Figures 4 and 5, respectively. F: female. M: male. Detailed cross results are found in Table S2. Crosses may have variable results depending on the replicate experiment. In crosses after a new thaw of the *C. ubi* and *C*. sp. 41 strains, the fertile F1 males gave rise to mostly males when crossed to *C*. sp. 41 females and to partially fertile males and females when crossed to *C. ubi f*emales, which could allow a population to propagate.

### Species declarations

The electronic edition of this article conforms to the requirements of the amended International Code of Zoological Nomenclature, and hence the new names contained herein are available under that Code from the electronic edition of this article. This published work and the nomenclatural acts it contains have been registered in ZooBank, the online registration system for the ICZN. The ZooBank LSIDs (Life Science Identifiers) can be resolved and the associated information viewed through any standard web browser by appending the LSID to the prefix ‘‘http://zoobank.org/’‘. The LSID for this publication is: urn:lsid:zoobank.org:pub:68091B60-E71A-4196-92E7-6033E400AD8C. The electronic edition of this work was published in a journal with an ISSN.

Most *Caenorhabditis* species in the *Elegans* group are highly similar morphologically, therefore we define them solely based on the biological species concept. Crosses were prioritized using genetic proximity (Félix et al. 2014). We provide pictures of their male tail in Figure S2 to conform with the ICZN guidelines.

#### *Caenorhabditis indonesiana* Tarno and Félix sp. n

Zoobank identifier

urn:lsid:zoobank.org:act:6D761875-0EC7-45D6-AD06-749DB4A0B124

= *Caenorhabditis* sp. 77 (temporary number)

The type isolate by present designation is HPT10. This strain is derived from a single female. The strain thus includes a single species. The holotype is deposited at the *Caenorhabditis* Genetics Center under the strain name HPT10 and paratypes are deposited as frozen strains at the Museum Koenig Bonn under ID#ZFMK-TIS-99298 to #ZFMK-TIS-99303. The species reproduces through males and females. The species is delineated and diagnosed by the fertile cross with the type isolate HPT10 in both cross directions, yielding highly fertile hybrid females and males that are interfertile and cross-fertile with their parent strains. This species differs by ITS2 DNA sequence from all species listed in Tables 1 and 2 of (Félix et al. 2014), those described in (Huang et al. 2014; Ferrari et al. 2017; Slos et al. 2017; Kanzaki et al. 2018; Crombie et al. 2019; Stevens et al. 2019; Dayi et al. 2021; Sloat et al. 2022) and other species in Table 2 of the present article. Note that these ribosomal DNA sequences may vary among repeated copies and within the species. From RNA sequence data, the closest species is *C. wallacei*, with which it does not form any adult progeny (Table S2). The type isolate was collected from a rotting banana flower collected in a forest near Batu, East Java, Indonesia (GPS -7.803387, 112.516604) on 28 April 2024. Other isolates were found in decomposing vegetal matter in East Java, Lombok and South Sulawesi, Indonesia (Table S1). The fan of the male tail is wide (Figure S2). Dorsal rays are found in antero-posterior positions 5 and 7. The males mate in a parallel position. The species is named after its place of isolation.

**Table 2.** New *Caenorhabditis* species.

| Species | Temporary number | Type strain | Mode of reproduction | 3-letter abbreviation | RNAseq accession number | Transcriptome assembly accession |
| --- | --- | --- | --- | --- | --- | --- |
| *indonesiana* | 77 | HPT10 | Male-female | Cid | SAMN48179281 | GLGD00000000 |
| *malinoi* | 78 | HPT35 | Male-female | Cml | SAMN48179282 | GLGE00000000 |
| *ceno* | 79 | HPT43 | Male-female | Cce | SAMN48179283 | GLGF00000000 |
| *brawijaya* | 80 | HPT50 | Male-female | Cbw | SAMN48179284 | GLGG00000000 |
| *ubi* | 81 | HPT5 | Male-female | Cub | SAMN48179285 | GLGC00000000 |

*indonesiana*: Indonesian. *malinoi*: of Malino. *ceno*: from the mud (Latin). *brawijaya*: Java prince and name of the university. *ubi*: for Universitas Brawijaya (UB). 3-letter abbreviations avoid repeating that used for other *Caenorhabditis* species. Alternatively, 4 letters can be used.

#### *Caenorhabditis malinoi* Tarno and Félix sp. n

Zoobank identifier

urn:lsid:zoobank.org:pub:68091B60-E71A-4196-92E7-6033E400AD8C

= *Caenorhabditis* sp. 78 (temporary number)

The type isolate by present designation is HPT35. This strain is derived from a single female. The strain thus includes a single species. The holotype is deposited at the *Caenorhabditis* Genetics Center under the strain name HPT35 and paratypes are deposited as frozen strains at the Museum Koenig Bonn under ID#ZFMK-TIS-99304 to #ZFMK-TIS-99309. The species reproduces through males and females. The species is delineated and diagnosed by the fertile cross with the type isolate HPT35 in both cross directions, yielding highly fertile hybrid females and males that are interfertile and cross-fertile with their parent strains. This species differs by ITS2 DNA sequence from all species listed in Tables 1 and 2 of (Félix et al. 2014), those described in (Huang et al. 2014; Ferrari et al. 2017; Slos et al. 2017; Kanzaki et al. 2018; Crombie et al. 2019; Stevens et al. 2019; Dayi et al. 2021; Sloat et al. 2022) and other species in Table 2 of the present article. Note that these ribosomal DNA sequences may vary among repeated copies and within the species. From RNA sequence data, the closest species are those in the clade formed by *C. zanzibari, C. tribulationis, C. sinica, C*. sp. 41 and the species described in the present article as *C. ceno, C. brawijaya* and *C. ubi*. The closest species is *C. ceno*. It does not form fertile hybrids with any of these species. *C. malinoi* females crossed to *C. ceno* males produce sterile males (Table S2). The type isolate was collected from rotting flowers of *Stewartia pseudocamellia* collected in a forest park in Malino, South Sulawesi, Indonesia (GPS -5.242944, 119.868592) on 6 May 2024. Other isolates were found in rotting fruits in other locations around Malino (Table S1). The fan of the male tail is wide (Figure S2). Dorsal rays are found in antero-posterior positions 5 and 7. The anterior side of the hook shows a distinctive three-lobed shape, which is a shared character with *C. sinica* (Huang et al. 2014), *C. tribulationis* and *C. zanzibari* (Stevens et al. 2019). The males mate in a parallel position. The species is named after its place of isolation.

#### *Caenorhabditis ceno* Tarno and Félix sp. n

Zoobank identifier

urn:lsid:zoobank.org:act:C3A6D80C-9B1F-45A1-A2A4-38FC8A38F132

= *Caenorhabditis* sp. 79 (temporary number)

The type isolate by present designation is HPT43. This strain is derived from a single female. The strain thus includes a single species. The holotype is deposited at the *Caenorhabditis* Genetics Center under the strain name HPT43 and paratypes are deposited as frozen strains at the Museum Koenig Bonn under ID#ZFMK-TIS-99310 to #ZFMK-TIS-99314. The species reproduces through males and females. The species is delineated and diagnosed by the fertile cross with the type isolate HPT43 in both cross directions, yielding highly fertile hybrid females and males that are interfertile and cross-fertile with their parent strains. This species differs by ITS2 DNA sequence from all species listed in Tables 1 and 2 of (Félix et al. 2014), those described in (Huang et al. 2014; Ferrari et al. 2017; Slos et al. 2017; Kanzaki et al. 2018; Crombie et al. 2019; Stevens et al. 2019; Dayi et al. 2021; Sloat et al. 2022) and other species in Table 2 of the present article. Note that these ribosomal DNA sequences may vary among repeated copies and within the species. From RNA sequence data, the closest species are those in the clade formed by *C. zanzibari, C. tribulationis, C. sinica, C*. sp. 41 and the species described in the present article as *C. malinoi, C. brawijaya* and *C. ubi*. The closest species is *C. malinoi*. It does not form fertile hybrids with any of these species. *C. ceno* males crossed to *C. malinoi* females produce sterile males. *C. ceno* females crossed to *C. ubi* males produce small hybrid males and to *C. tribulationis* males produce small hybrid females (Table S2). The type isolate was collected from leaf litter collected in a forest in South Sulawesi, Indonesia (GPS - 5.238297, 119.642281) on 6 May 2024. Two other isolates were found in the same forest (Table S1). The fan of the male tail is wide (Figure S2). Dorsal rays are found in antero-posterior positions 5 and 7. The anterior side of the hook shows a distinctive three-lobed shape that is shared with *C. sinica* (Huang et al. 2014), *C. tribulationis* and *C. zanzibari* (Stevens et al. 2019). The males mate in a parallel position. The species is named after the mud in the forest where it was collected (from the Latin cenum).

#### *Caenorhabditis brawijaya* Tarno and Félix sp. n

Zoobank identifier

urn:lsid:zoobank.org:act:3058D518-A8DA-44DA-90FC-DD826A5D19A7

= *Caenorhabditis* sp. 80 (temporary number)

The type isolate by present designation is HPT50. This strain is derived from a single female. The strain thus includes a single species. The holotype is deposited at the *Caenorhabditis* Genetics Center under the strain name HPT50 and paratypes are deposited as frozen strains at the Museum Koenig Bonn under ID#ZFMK-TIS-99315 to #ZFMK-TIS-99319. The species reproduces through males and females. The species is delineated and diagnosed by the fertile cross with the type isolate HPT50 in both cross directions, yielding highly fertile hybrid females and males that are interfertile and cross-fertile with their parent strains. This species differs by ITS2 DNA sequence from all species listed in Tables 1 and 2 of (Félix et al. 2014), those described in (Huang et al. 2014; Ferrari et al. 2017; Slos et al. 2017; Kanzaki et al. 2018; Crombie et al. 2019; Stevens et al. 2019; Dayi et al. 2021; Sloat et al. 2022) and other species in Table 2 of the present article. Note that these ribosomal DNA sequences may vary among repeated copies and within the species. From RNA sequence data, the closest species are the *C. zanzibari, C. tribulationis, C. sinica, C*. sp. 41 and the species described in the present article as *C. malinoi, C. ceno* and *C. ubi*. The closest and present sister species is *C. sinica. C. brawijaya* does not form fertile hybrids with any of these species but males crossed to *C. sinica* yielded some adult female progeny and females crossed to *C. zanzibari* males produce some sterile males (Table S2). The type isolate was collected from a rotting *Musa* pseudostem collected in a forest near Ngadas, Malang Regency, East Java, Indonesia (GPS -7.99746, 112.87387) on 11 May 2024. Another isolate was found in decomposing leaves a few kilometers away (Table S1). The fan of the male tail is wide (Figure S2). Dorsal rays are found in antero-posterior positions 5 and 7. The anterior side of the hook shows a distinctive three-lobed shape that is shared with *C. sinica* (Huang et al. 2014), *C. tribulationis* and *C. zanzibari* (Stevens et al. 2019). The males mate in a parallel position. The species is named after the Javanese prince after whom the Universitas Brawijaya is named.

#### *Caenorhabditis ubi* Tarno and Félix sp. n

Zoobank identifier

urn:lsid:zoobank.org:act:2B4292DE-2F01-4E87-8E11-E82811024443

= *Caenorhabditis* sp. 81 (temporary number)

The type isolate by present designation is HPT5, deposited at the Caenorhabditis Genetics Center. This strain is derived from a single female. The strain thus includes a single species. The holotype is deposited at the *Caenorhabditis* Genetics Center under the strain name HPT5 and paratypes are deposited as frozen strains at the Museum Koenig Bonn under ID#ZFMK-TIS-99320 to #ZFMK-TIS-99324. The species reproduces through males and females. The species is delineated and diagnosed by the fertile cross with the type isolate HPT5 in both cross directions, yielding highly fertile hybrid females and males that are interfertile and cross-fertile with their parent strains. This species differs by ITS2 DNA sequence from all species listed in Tables 1 and 2 of (Félix et al. 2014), those described in (Huang et al. 2014; Ferrari et al. 2017; Slos et al. 2017; Kanzaki et al. 2018; Crombie et al. 2019; Stevens et al. 2019; Dayi et al. 2021; Sloat et al. 2022) and other species in Table 2 of the present article. Note that these ribosomal DNA sequences may vary among repeated copies and within the species. From RNA sequence data, the closest species are *C*. sp. 41, *C. zanzibari, C. tribulationis, C. sinica*, and the species described in the present article as *C. malinoi, C. ceno* and *C. brawijaya*. It forms some fertile hybrids in some cross directions with *C*. sp. 41 as described in the present article (Table S2, Figure 2). The type isolate was collected from a rotting banana flower collected near Batu, East Java, Indonesia (GPS -7.803387, 112.516604) on 28 April 2024 (Table S1). The fan of the male tail is wide (Figure S2). Dorsal rays are found in antero-posterior positions 5 and 7. The anterior side of the hook shows a distinctive three-lobed shape that is shared with *C. sinica* (Huang et al. 2014), *C. tribulationis* and *C. zanzibari* (Stevens et al. 2019). The males mate in a parallel position. The species is named after the Universitas Brawijaya (UB).

### Phylogenetic relationships including closely related pairs of species

From our phylogenetic reconstruction using RNA sequencing of 1,861 orthologous single-copy genes (Figure 2), four of these new species belong to a *Sinica* subclade of species so far only found in an East Asia-Indo-Pacific world region. This clade includes two pairs of particularly closely related species with short branches on the tree and low distances (Figure 2, Table S4): *C. ubi* with *C*. sp. 41 from the Solomon islands, and *C. ceno* with *C. malinoi*, the latter two both from 30 km apart in South Sulawesi.

The fifth species, *C. indonesiana*, appears as the sister of the *C. tropicalis*-*C. wallacei* pair. *C. tropicalis* was found in our collection in Indonesia and is cosmopolitan in tropical regions. *C. wallacei* was previously found in Bali, Indonesia. The outgroup is *C. doughertyi*, found in South India (Kiontke et al. 2011; Félix et al. 2014).

### Hybrid crosses that break Haldane’s rule

Haldane’s rule regarding species hybrids says that individuals of the heterogametic sex are the first affected in hybrid crosses (Haldane 1922), whether by lethality, slow growth or sterility. Asymmetric results may in addition be obtained between the two cross directions (Darwin’s “corollary” to Haldane’s rule). In *Caenorhabditis*, males are the heterogametic sex (X0), thus according to Haldane’s rule are more likely to be affected than the females. Accordingly, a pattern observed in many *Caenorhabditis* species crosses is that only hybrid F1 females are produced in the progeny, as observed in (Baird et al. 1992; Woodruff et al. 2010; Kiontke et al. 2011; Kozlowska et al. 2012; Bi et al. 2019). We observed an asymmetric pattern corresponding to Haldane’s rule with an excess of females or slow-growing and small males in crosses between *C. ceno* females and *C. tribulationis* males (Figure 3C) or *C*. sp. 41 females and *C. ubi* males (Figure 3D).

However, other crosses instead yielded males only, for example the *C. malinoi* females crossed with *C. ceno* males, *C. ceno* females crossed to *C. ubi* males, or *C. tribulationis* females with *C. ubi* males (Table S2, Figures 2A,B and 3). In some cases, the males were abnormally small (Figure 4). We focused on the new species pair *C. malinoi* and *C. ceno*, collected a few kilometers apart in South Sulawesi. The result was consistent when multiple strains of *C. malinoi* and *C. ceno* were tested (Table S2). In the case of these two species, the numerous and normal-size F1 males could not sire progeny and did not deposit a mating plug on females. When observed in the compound microscope, many individuals showed a defective connection between the gonad and the rectum (Figure 4).

**Figure 4.**
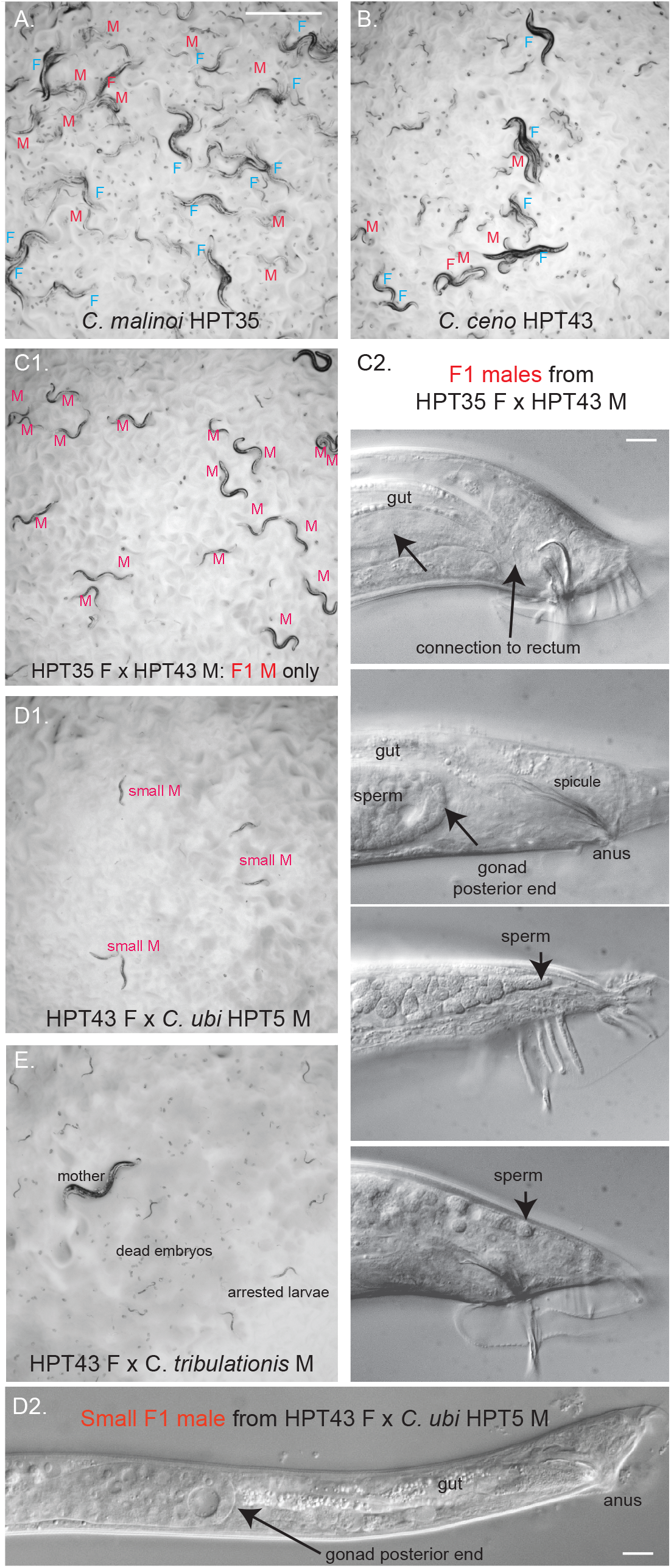
Hybrid crosses between *Caenorhabditis* species break Haldane’s rule. Crosses between the two *Caenorhabditis* species from South Sulawesi, *C. malinoi* and *C. ceno* (A,B), yield males only (C), thus breaking Haldane’s rule concerning viability. (C2) These males are however sterile and display a defective connection between the gonad and the cloaca (rectum), thus preventing the transfer of sperm and of the mating plug material. The (C2) panel shows Nomarski micrographs of F1 hybrid males: on top an apparently normal male and below three adult males with a defective gonad-cloaca connection. (D) shows the adult males of small size obtained in the cross of *C. ceno* HPT43 females to *C. ubi* HPT5 males. (D1) shows them on the plate at the same magnification as (A-C). (D2) shows a Nomarski micrograph at the same scale as (C2). (E) shows an example of a cross yielding only arrested larvae and many dead embryos, in this case *C. ceno* HPT43 females crossed with *C. tribulationis* JU2774 males. Panels A, B, C1, D1, E have the same magnification and the corresponding scale bar in panel A represents 1 mm. Bars in panel C2 and D2: 10 μm.

### Partial fertility of hybrids between *C*. sp. 41 and *C. ubi*

Studying genetic incompatibility using genetic mapping is possible when the first generation (F1) progeny can produce second generation (F2) progeny, in which reassortment of chromosomes and recombination may have occurred. We found such a case here with *C. ubi* from a forest in East Java and *C*. sp. 41 from Solomon Islands.

The cross of *C*. sp. 41 BRC20276 females with *C. ubi* HPT5 males - let’s call this direction 41ubi - gave rise mostly to F1 females, following Haldane’s rule (Figure 2D, left). These 41ubi F1 females did not give rise to any progeny when placed with males of either parental strain.

By contrast, reverse crosses between *C. ubi* HPT5 females and *C*. sp. 41 BRC20276 males (ubi41) yielded partially fertile F1 adults of both sexes, with the males being most fertile (Figure 2D, right). When crossed to each other, these F1 hybrids yielded few abnormal F2 progeny and rarely could the population propagate. The F1 hybrid females crossed with *C*. sp. 41 BRC20276 males did not give progeny but yielded some progeny when backcrossed to *C. ubi* HPT5 males (Figure 5). In several replicate experiments (e.g. Figure 5), when crossed to HPT5 females, the F1 hybrid males gave rise to few small and sterile adult progeny of both sexes and, most interestingly, when crossed to BRC20276 females, could yield fertile progeny. The lineages from could be continued for several generations. These crosses also gave rise to many dead embryos and arrested larvae, including what appeared to be a predauer or dauer-like stage.

**Figure 5.**
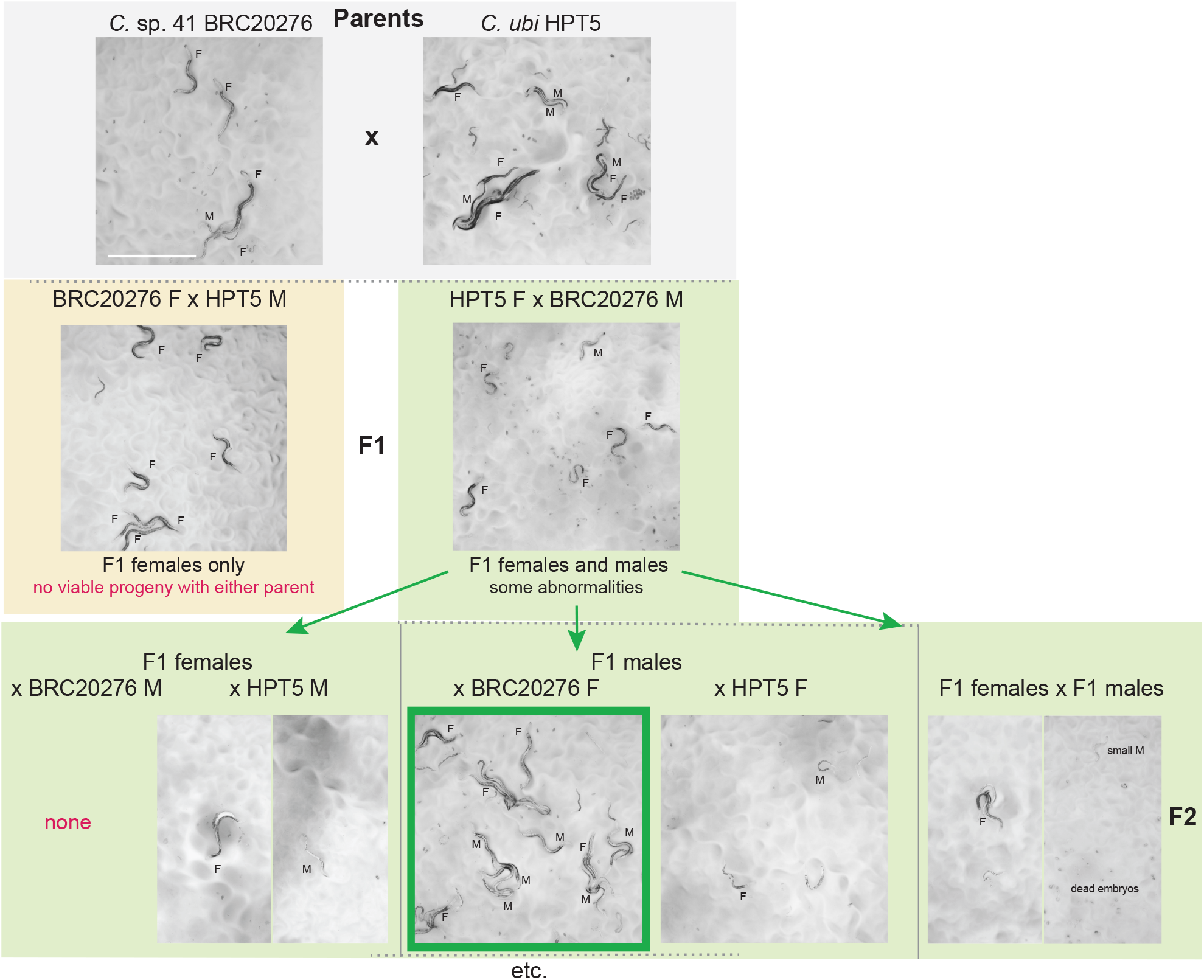
Pictures of the progeny of crosses between *C. ubi* HPT5 and *C*. sp. 41 BRC20276, yielding partially fertile hybrids. Animals of the parental, F1, F2 and backcrosses are shown. In this experiment, only the backcross of the F1 hybrid males with BRC20276 females yielded abundant fertile progeny. The crosses are schematized in Figure 2D. F: female. M: male. Bar: 1 mm. Same magnification for all panels. Note that the results of the backcrosses with F1 males can yield different results due to within-strain polymorphisms in incompatibility loci in the parental strains.

The parental strains are isofemale lines but not isogenic. In experiments performed on a new thaw of the strains, the backcross of ubi41 F1 males to HPT5 females gave abundant progeny of both sexes, while that to BRC20276 mostly gave rise to males (2 replicates each). The difference with the result shown in Figures 3 and 5 may be explained by polymorphisms for incompatibility segregating in one or both parental strains. In any case, the initial cross using *C. ubi* females and *C*. sp. 41 males followed by a backcross of ubi41 F1 males appears to be a good strategy to produce hybrids between the two species.

Thus, *C. ubi* and *C*. sp. 41 are distinct biological species but can yield fertile lineages. This partial reproductive isolation is ideal for studies of incompatibility between closely related species and thus studies of genetic and molecular mechanisms of speciation.

## Discussion

### Biodiversity and biogeography

The present study found eleven different *Caenorhabditis* species in Indonesia, ten of which could be kept in culture and frozen, forming a collection of 62 strains. Indonesia is rich in species of the *Elegans* group: all collected species were from this group and represent half of the known *Elegans* group species (Figure 2). Of the five previously found species, all occur on several continents, including the three selfing species.

Five species of the *Elegans* group are newly described here. *C. indonesiana* is a close relative of *C. wallacei* and *C. tropicalis* and has been found on three Indonesian islands. The four others were each found on a single island and belong to the *Sinica* subclade of the *Elegans* goup that includes *C. sinica, C. zanzibari, C. tribulationis*, and *C*. sp. 41. This *Sinica* subclade has been found in China (*C. sinica*), Australia (*C. tribulationis*) Solomon islands (*C*. sp. 41) and islands East of Africa in the Indian ocean (*C. zanzibari*). Besides geography, this species group is characterized by the trilobed shape of the male sensory organ located anterior to the cloaca called the hook (Kiontke et al. 2011; Huang et al. 2014; Stevens et al. 2019) (see Figure S2 legend). Altogether, Indonesia appears to harbor species belonging to several subclades of the *Elegans* group and many potentially endemic species, particularly in two subclades. This is consistent with the possibility that this region of the world having been a source of diversification of the *Elegans* group of *Caenorhabditis* species (Kiontke et al. 2011; Frézal and Félix 2015).

*C. elegans* seems to fare less well than others at high temperatures in the laboratory (Félix and Duveau 2012; Begasse et al. 2015; Frézal et al. 2023) and in the field (Kiontke et al. 2011; Crombie et al. 2019). It was found here only once at a high elevation (2,156 meters). Consistent with their sampling at higher elevation (1,500-2,400 meters), cultures of *C. elegans* HPT48 and *C. brawijaya* HPT49 and HTP50 cannot grow beyong a few generations at 25°C on *E. coli* OP50 while they thrive at 20°C.

### Violations of Haldane’s rule

Our crosses of species within the *Sinica* subclade yielded interesting patterns in the hybrids, the first one being the preferential occurrence of F1 males in some crosses, another providing a pair of species that can serve for genetic studies of incompatibilities arising between closely related species.

Haldane’s “rule” corresponds to an overwhelming pattern in species hybrids across organisms, where the heterogametic sex is the first affected in terms of lethality, growth or sterility. In *Caenorhabditis* species where males carry a single X chromosome, one mechanism explaining the Haldane pattern is an incompatibility between the maternal X chromosome in the hemizygous state (single copy in X0 males) and the rest of the genome in the heterozygous state, while it is rescued by the second X chromosome of the other parent in females (Laurie 1997; Cutter 2024; Li et al. 2024). In a recently studied case between *C. briggsae* and *C. nigoni, xol-1*, a key gene in sex determination and dosage compensation being located on the X chromosome, is insufficiently expressed in sterile hybrid males (Li et al. 2024). An alternative or more extreme mechanism for Haldane’s rule is sexual transformation of these X0 animals in females (Baird 2002). Additional mechanisms may be at play (Laurie 1997; Cutter 2024).

Any biological “rule” suffers exceptions. Here we find crosses that contradict the Haldane pattern and only yield hybrid males. Such examples have been found in insects (Laurie 1997); for example in fruitflies, a cross of *Drosophila simulans* females with *D. melanogaster* males results in hybrid males dying as larvae while females die as embryos (Sawamura et al. 1993). In this case, a mismatch between the mother’s cytoplasmic content and the mitotic segregation of the father’s X chromosome in the early embryo is responsible for the embryonic lethality of females (Ferree and Barbash 2009). Several mechanisms may in principle explain a violation of Haldane’s rule: i) as in this *Drosophila* example, an incompatibility between the paternal X chromosome and the maternal autosomes or cytoplasmic content (e.g. mitochondria, small RNAs, maternal content of proteins), preventing development or fertility of females; ii) incompatibility between the two X chromosomes, masculinization of XX animals and/or a defect in dosage compensation of XX females; iii) in nematodes with X0 males, loss of one X chromosome either by selection of sperm not bearing it or by mispairing in oogenesis after fertilization. In addition, we cannot rule out a novel sex determination system, for example ZZ/ZW (Hodgkin 2002).

In the examples found here in *Caenorhabditis*, the reverse crosses did not yield any adult animals (Figure 2), which is compatible with several of the above scenarios, including the paternal X-autosome incompatibility if the paternal X acts dominantly in females. In addition to the X chromosome, mitochondria and sex determination loci are both likely to evolve fast in these species and are thus prime candidates for such “non-Haldanian” patterns. We note that the two crosses yielding males have a *C. ceno* parent but as males with *C. malinoi* females, and as females with *C. ubi* males. This is consistent with the possibility of a defective dosage compensation in females in the presence of an X chromosome from *C. ceno*, instead of being aberrantly activated in males as in the *C. briggsae* - *C. nigoni* case (Li et al. 2024). Further studies aimed at understanding this exception to Haldane’s rule may particularly benefit from the *C. ceno-C. malinoi* pair of sister species, isolated 30 kilometers from each other and yielding many males of normal size.

### Gonad-proctodeum connection as a weak point in male development?

The *C. ceno-C. malinoi* cross reproduces a defect previously seen in *C. remanei*-*C. latens* crosses, where in one cross direction the F1 males are sterile, and the male sterility is explained by a defective gonad-cloaca connection (Dey et al. 2014). We observed such defects in F1 males after crossing *C. ceno* males to *C. ubi* males (Figure 3) as well as rare F1 males obtained by crossing *C. tribulationis* females to *C. ubi* males (Table S2). In the latter case, we observed that the gut connection to the rectum was also defective, resulting in constipated small animals that could not feed normally. *C. remanei* and *C. latens* are not particularly closely related t*o* these *Sinica* subgroup species. We thus propose that gonad connection driven by the migration of the somatic linker cell may be a weak point in *Caenorhabditis* male development. Mutants defective in this connection have been found in *C. elegans* (Palmer et al. 2002; Kato and Sternberg 2009; Kato et al. 2021; Smith et al. 2024). A possibility is that this defective gonad-proctodeum results in multiple species from aberrant dosage compensation in males or a partial sexual transformation affecting this developmental trait more than others.

### A pair of partially cross-fertile species

*C. ubi* and *C*. sp. 41 provide a new case of partially fertile hybrids in the genus, with a pattern distinct from the pairs *C. briggsae* - *C. nigoni* and *C. remanei* - *C. latens*. Here the successful cross at the second generation in the experiments shown in Figures 3 and 5 is a backcross of F1 males, which can further produce progeny. This cross-fertility opens the door to further genetic tests and recombinant mapping. Two other backcrosses yielded none or far fewer progeny (Figure 5), however they could also be propagated after a bottleneck.

The most restrictive parental cross (41ubi) gave rise mostly to F1 females, abiding by Haldane’s rule. The cross of *C. ubi* females with *C*. sp. 41 males (ubi41) yields both females and males. The ubi41 F1 females could be mated, produce embryos and a few adult progeny with the maternal *C. ubi* parent. By contrast, the ubi41 F1 males appeared more fertile than the females. This can be considered to contradict Haldane’s rule in terms of sterility of these ubi41 hybrids; arguably, it may or may not correspond to a physiological sterility of the F1 females since they form F2 embryos, which generally arrest.

This pattern may suggest a X chromosome effect where the *C*. sp. 41 (X41) chromosome in the hemizygous state is incompatible with *C. ubi* autosomes, producing the lethality of the 41ubi F1 males bearing a single X41. Other explanations are possible, such as feminization of hybrid X0 animals bearing the X41 chromosome (Baird 2002; Cutter 2024). A X41 chromosome effect may also explain the greater fertility of the ubi41 F1 males with a single X chromosome from *C. ubi*, compared to their sisters that carry a X chromosome from both parental species. In this case, the X41 would act dominantly on fertility in females. The difference between crosses of *C*. sp. 41 females to *C. ubi* versus to ubi41 F1 males (both with the *C. ubi* X chromosome), where males carrying a hemizygous X41 develop in the latter case only, could be explained by the difference in autosomal content.

What cannot be explained with a X41-autosome incompatibility is that the ubi41 F1 males with Xubi are less cross-fertile with *C. ubi* females than with *C*. sp. 41 females, at least in the series of experiments shown in Figures 3 and 5. A simple mitochondrial or maternal effect is not an obvious explanation since their mother is *C. ubi*. A paternal effect of the *C*. sp. 41 fathers could be involved. Multiple incompatibilites and/or sex transformations may coexist, resulting in the observed pattern. In addition, as mentioned above, the parental strains are not isogenic and likely segregate genetic variation for these incompatibilities. Further genetic studies with genotyping will be required to solve this genetic puzzle and address genetic incompatibility mechanisms.

## Data availability

Strains are available upon request. The ribosomal DNA ITS2 sequences are deposited at GenBank with accession numbers PV569245-PV569249. Sequence reads and transcriptome assemblies are available at NCBI with accession numbers listed in Table 2.

## Acknowledgements

We thank Charlie Baer, Shu-Dan Yeh and Anggun Sausan Firdaus for establishing the connection between the authors. We thank Hery Haryanto and L. Sukardi for help and participation to sample collection. We thank Luqman Qurata Aini for help with plate pouring and bacterial culture, Xu Wei for help with coordinate mapping, Marie Delattre for discussion. This work was funded by the Centre National de la Recherche Scientifique (France), Universitas Brawijaya (Indonesia), and the United States National Institute of General Medical Sciences (GM141906).

## Figure legends

## Supplemental Information

**Figure S1. Location and sample yielding *C. elegans***. For the *C. elegans* isolate, the pictures show the sampling site location at the bottom of the Batok and Bromo volcanoes, the decomposing stems and the laboratory culture (HPT48).

**Figure S2. Male tail micrographs of the five newly described species**. Nomarski micrographs of individuals of the reference strain. Left: left lateral view. Right: Ventral view. Two different nomenclatures for tail sensory organs are used, one counting nine rays from anterior to posterior, the other distinguishing the ventral from the dorsal ones. In all species, the anterior dorsal ray (ad) is the fifth and the posterior dorsal ray (pd) is the seventh. The precloacal sensillum has a hook shape in all species, and is trilobed at least in all four species of the *Sinica* subclade. We note here that the lateral lobes of this precloacal sensory organ appear to be slightly more dorsal than the central part and could derive from fusion of the cuticle (dorso)-lateral to the central part of the hook (see arrowhead on the *C. ceno* HPT43 ventral view, for which the focal plane was chosen to demonstrate this). We further note that *C. indonesiana* HPT10, outside of the *Sinica* subclade, has a hint of a trilobed hook as well, with the cuticle dorso-lateral to the central part of the hook tending to attach to it. Bar: 10 μm, valid for all panels.

**Table S1. Samples and strains**. Each sheet corresponds to a sampling location. Each line is a sample, and those positive for *Caenorhabditis* are indicated. Some samples produced two or three different *Caenorhabditis* species. The data are summarized in Table 1.

**Table S2. Mating tests**. This table shows the result of crossing tests using five L4 stage females and five males. The presence and developmental stage of progeny were assessed over several days. Each tested cell of the matrix is color-coded according to the most advanced developmental stage of the hybrid progeny, as indicated at the bottom of the table. The number indicates the number of independent mating tests, for example 2/2 indicates 2 crosses with the same result.

**Table S3. Links to sequencing data used for the phylogenetic tree**.

**Table S4. Pairwise distance matrix along branches of the phylogenetic tree**.

## References

Baird, S. E. (2002). Haldane’s rule by sexual transformation in Caenorhabditis. Genetics 161: 1349–1353.

Baird, S. E., Sutherlin, M. E., Emmons, S. W. (1992). Reproductive isolation in Rhabditidae (Nematoda, Secernentea) - mechanisms that isolate six species of three genera. Evolution 46: 585–594.

Barrière, A., Félix, M.-A. (2014). Isolation of C. elegans and related nematodes. Wormbook DOI: 10.1895/wormbook.1.115.2.

Begasse, M. L., Leaver, M., Vazquez, F., Grill, S. W., Hyman, A. A. (2015). Temperature dependence of cell division timing accounts for a shift in the thermal limits of C. elegans and C. briggsae. Cell Rep 10: 647–653.

Bi, Y., Ren, X., et al. (2019). Specific interactions between autosome and X chromosomes cause hybrid male sterility in Caenorhabditis species. Genetics 212: 801–813.

Bi, Y., Ren, X., et al. (2015). A genome-wide hybrid incompatibility landscape between Caenorhabditis briggsae and C. nigoni. PLoS Genet 11: e1004993.

Bundus, J. D., Alaei, R., Cutter, A. D. (2015). Gametic selection, developmental trajectories, and extrinsic heterogeneity in Haldane’s rule. Evolution 69: 2005–17.

Capella-Gutierrez, S., Silla-Martinez, J. M., Gabaldon, T. (2009). trimAl: a tool for automated alignment trimming in large-scale phylogenetic analyses. Bioinformatics 25: 1972–3.

Crombie, T. A., McKeown, R., et al. (2023). CaeNDR, the Caenorhabditis Natural Diversity Resource. Nucleic Acids Res 52: D850–D858.

Crombie, T. A., Zdraljevic, S., et al. (2019). Deep sampling of Hawaiian Caenorhabditis elegans reveals high genetic diversity and admixture with global populations. Elife 8: e50465.

Cutter, A. D. (2017). X exceptionalism in Caenorhabditis speciation. Mol Ecol 27: 3925–3934.

Cutter, A. D. (2024). Beyond Haldane’s rule: Sex-biased hybrid dysfunction for all modes of sex determination. Elife 13: e96652.

Dayi, M., Kanzaki, N., et al. (2021). Additional description and genome analyses of Caenorhabditis auriculariae representing the basal lineage of genus Caenorhabditis. Sci Rep 11: 6720.

Dey, A., Jeon, Y., Wang, G. X., Cutter, A. D. (2012). Global population genetic structure of Caenorhabditis remanei reveals incipient speciation. Genetics 191: 1257–69.

Dey, A., Jin, Q., Chen, Y. C., Cutter, A. D. (2014). Gonad morphogenesis defects drive hybrid male sterility in asymmetric hybrid breakdown of Caenorhabditis nematodes. Evol Dev 16: 362–72.

Félix, M.-A., Braendle, C. (2010). The natural history of Caenorhabditis elegans. Curr Biol 20: R965–9.

Félix, M. A., Braendle, C., Cutter, A. D. (2014). A streamlined system for species diagnosis in Caenorhabditis (Nematoda: Rhabditidae) with name designations for 15 distinct biological species. PLoS One 9: e94723.

Félix, M. A., Duveau, F. (2012). Population dynamics and habitat sharing of natural populations of Caenorhabditis elegans and C. briggsae. BMC Biol 10: 59.

Ferrari, C., Salle, R., et al. (2017). Ephemeral-habitat colonization and neotropical species richness of Caenorhabditis nematodes. BMC Ecol 17: 43.

Ferree, P. M., Barbash, D. A. (2009). Species-specific heterochromatin prevents mitotic chromosome segregation to cause hybrid lethality in Drosophila. PLoS Biol 7: e1000234.

Frézal, L., Félix, M. A. (2015). C. elegans outside the Petri dish. Elife 4: e05849.

Frézal, L., Saglio, M., et al. (2023). Genome-wide association and environmental suppression of the mortal germline phenotype of wild C. elegans. EMBO Rep: e58116.

Grabherr, M. G., Haas, B. J., et al. (2011). Full-length transcriptome assembly from RNA-Seq data without a reference genome. Nat Biotechnol 29: 644–52.

Haldane, J. B. S. (1922). Sex ratio and unisexual sterility in hybrid animals. J. Genet. 12: 101–109.

Hodgkin, J. (2002). Exploring the envelope. Systematic alteration in the sex-determination system of the nematode Caenorhabditis elegans. Genetics 162: 767–80.

Huang, R. E., Ren, X., Qiu, Y., Zhao, Z. (2014). Description of Caenorhabditis sinica sp. n. (Nematoda: Rhabditidae), a nematode species used in comparative biology for C. elegans. PLoS One 9: e110957.

Jauharlina Oktarina, H., et al. (2022). Association of fig pollinating wasps and fig nematodes inside male and female figs of a dioecious fig tree in Sumatra, Indonesia. Insects 13.

Kanzaki, N., Tsai, I. J., et al. (2018). Biology and genome of a newly discovered sibling species of Caenorhabditis elegans. Nat Commun 9: 3216.

Kato, M., Kolotuev, I., Cunha, A., Gharib, S., Sternberg, P. W. (2021). Extrasynaptic acetylcholine signaling through a muscarinic receptor regulates cell migration. Proc Natl Acad Sci U S A 118: e1904338118.

Kato, M., Sternberg, P. W. (2009). The C. elegans tailless/Tlx homolog nhr-67 regulates a stage-specific program of linker cell migration in male gonadogenesis. Development 136: 3907–15.

Katoh, K., Standley, D. M. (2013). MAFFT multiple sequence alignment software version 7: improvements in performance and usability. Mol Biol Evol 30: 772–80.

Kiontke, K., Félix, M.-A., et al. (2011). A phylogeny and molecular barcodes for Caenorhabditis, with numerous new species from rotting fruits. BMC Evol. Biol. 11: 339.

Kozlowska, J. L., Ahmad, A. R., Jahesh, E., Cutter, A. D. (2012). Genetic variation for postzygotic reproductive isolation between Caenorhabditis briggsae and Caenorhabditis sp. 9. Evolution 66: 1180–95.

Laurie, C. C. (1997). The weaker sex is heterogametic: 75 years of Haldane’s rule. Genetics 147: 937–51.

Lee, D., Zdraljevic, S., et al. (2021). Balancing selection maintains hyper-divergent haplotypes in Caenorhabditis elegans. Nat Ecol Evol 5: 794–807.

Li, Y., Gao, Y., et al. (2024). Regulatory divergences in dosage compensation cause hybrid male inviability in Caenorhabditis. bioRxiv: doi:10.1101/2024.02.06.577000

Nguyen, L. T., Schmidt, H. A., von Haeseler, A., Minh, B. Q. (2015). IQ-TREE: a fast and effective stochastic algorithm for estimating maximum-likelihood phylogenies. Mol Biol Evol 32: 268–74.

Palmer, R. E., Inoue, T., Sherwood, D. R., Jiang, L. I., Sternberg, P. W. (2002). Caenorhabditis elegans cog-1 locus encodes GTX/Nkx6.1 homeodomain proteins and regulates multiple aspects of reproductive system development. Dev Biol 252: 202–13.

Paradis, E., Schliep, K. (2019). ape 5.0: an environment for modern phylogenetics and evolutionary analyses in R. Bioinformatics 35: 526–528.

Sawamura, K., Yamamoto, M. T., Watanabe, T. K. (1993). Hybrid lethal systems in the Drosophila melanogaster species complex. II. The Zygotic hybrid rescue (Zhr) gene of D. melanogaster. Genetics 133: 307–13.

Schulenburg, H., Félix, M.-A. (2017). The natural biotic environment of Caenorhabditis elegans. Genetics 206: 55–86.

Seppey, M., Manni, M., Zdobnov, E. M. (2019). BUSCO: assessing genome assembly and annotation bompleteness. Methods Mol Biol 1962: 227–245.

Shaham, S. (2006). Methods in cell biology. WormBook DOI: 10.1895/wormbook.1.49.1.

Sloat, S. A., Noble, L. M., et al. (2022). Caenorhabditis nematodes colonize ephemeral resource patches in neotropical forests. Ecol Evol 12: e9124.

Slos, D., Sudhaus, W., Stevens, L., Bert, W., Blaxter, M. (2017). Caenorhabditis monodelphis n. sp.: defining the stem morphology and genomics of the genus Caenorhabditis. BMC Zoology 2: 4.

Smith, M., Lesperance, M., et al. (2024). Two C. elegans DM domain proteins, DMD-3 and MAB-3, function in late stages of male somatic gonad development. Dev Biol 514: 50–65.

Stevens, L., Félix, M.-A., et al. (2019). Comparative genomics of ten new Caenorhabditis species. Evolution Letters 3: 217–236.

Stiernagle, T. (2006). “Maintenance of C. elegans.” Wormbook, from https://www.ncbi.nlm.nih.gov/pubmed/18050451. DOI: 10.1895/wormbook.1.101.1.

Woodruff, G. C., Eke, O., Baird, S. E., Félix, M.-A., Haag, E. S. (2010). Fertile interspecies hybrids in Caenorhabditis nematodes and their implications for the evolution of hermaphroditism. Genetics 186: 997–1012.

Woodruff, G. C., Phillips, P. C. (2018). Field studies reveal a close relative of C. elegans thrives in the fresh figs of Ficus septica and disperses on its Ceratosolen pollinating wasps. BMC Ecol 18: 26.

Zhang, C., Rabiee, M., Sayyari, E., Mirarab, S. (2018). ASTRAL-III: polynomial time species tree reconstruction from partially resolved gene trees. BMC Bioinformatics 19: 153.

