## Supplementary figures and images for "Five new *Caenorhabditis* species from Indonesia provide exceptions to Haldane’s rule and partial fertility of interspecific hybrids"

### Fig S1

*C. elegans* HPT48

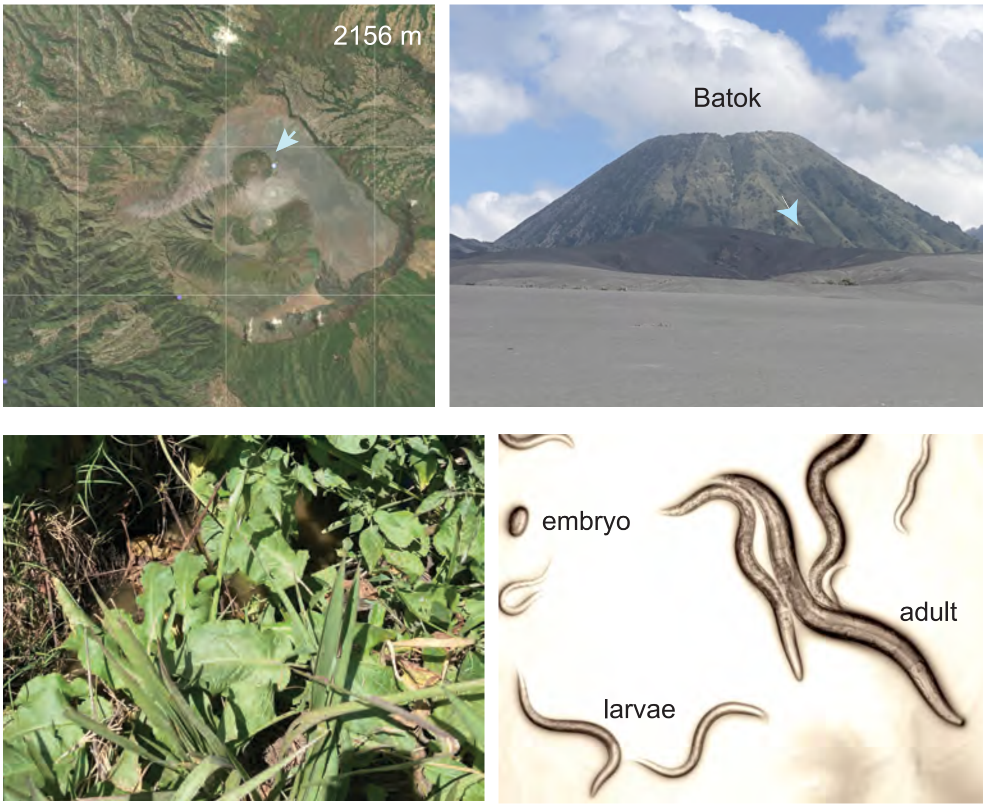


Figure S1

### Fig S2

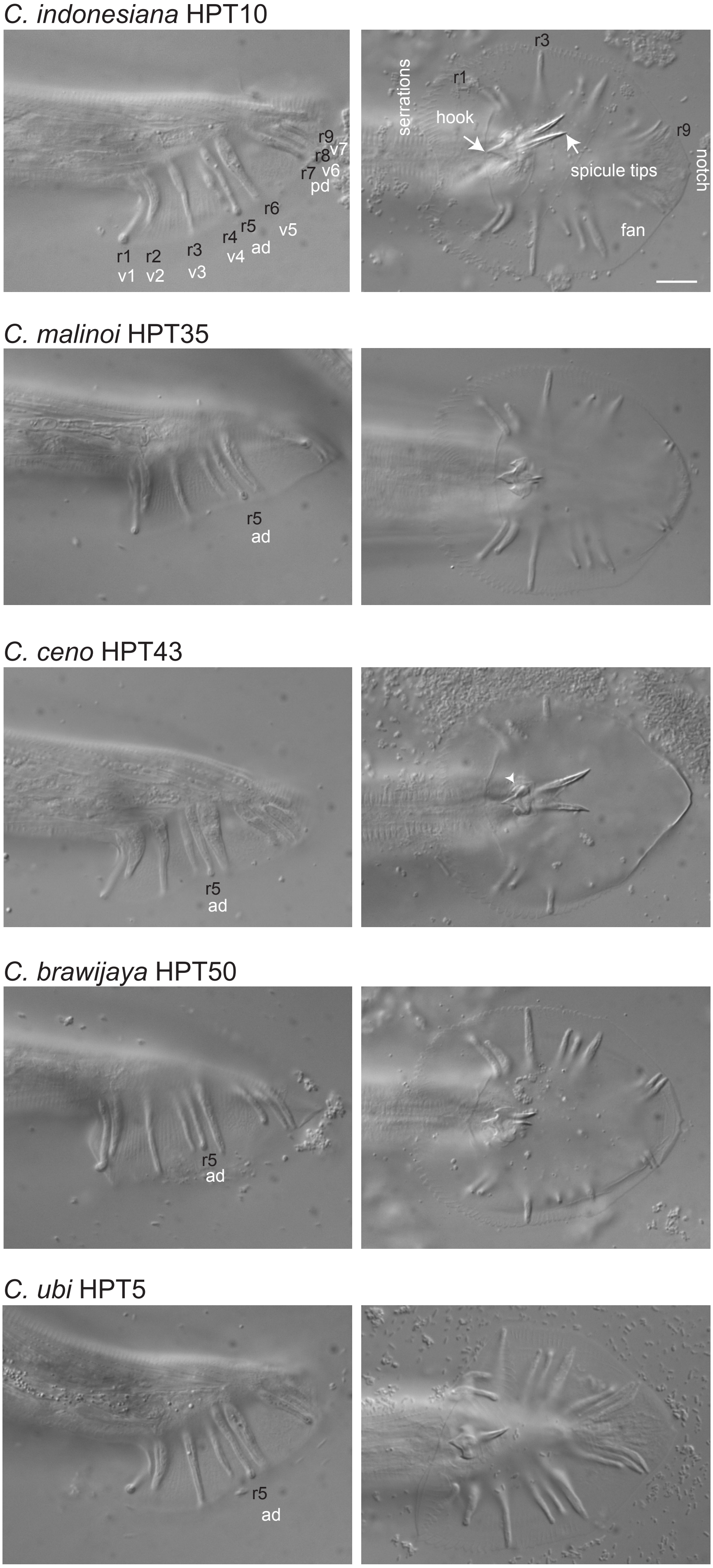
